# Disruption of nucleoid expanded conformation by toxic aberrant proteins synthesized in *Escherichia coli*

**DOI:** 10.1101/2021.10.06.463398

**Authors:** Mangala Tawde, Abdelaziz Bior, Michael Feiss, Paul Freimuth

## Abstract

Aminoglycoside antibiotics interfere with selection of cognate tRNAs during translation, resulting in the production of aberrant proteins that are the ultimate cause of the antibiotic bactericidal effect. To determine if these aberrant proteins are recognized as substrates by the cell’s protein quality control machinery, we studied whether the heat shock (HS) response was activated following exposure of *Escherichia coli* to the aminoglycoside kanamycin (Kan). Levels of the HS transcription factor σ32 increased about 10-fold after exposure to Kan, indicating that at least some aberrant proteins were recognized as substrates by the molecular chaperone DnaK. To investigate whether toxic aberrant proteins therefore might escape detection by the QC machinery, we studied model aberrant proteins that had a bactericidal effect when expressed in *E. coli* from cloned genes. As occurred following exposure to Kan, levels of σ32 were permanently elevated following expression of an acutely toxic 48-residue protein (ARF48), indicating that toxic activity and recognition by the QC machinery are not mutually exclusive properties of aberrant proteins, and that the HS response was blocked in these cells at some step downstream of σ32 stabilization. This block could result from halting of protein synthesis or from radial condensation of nucleoids, both of which occurred rapidly following ARF48 induction. Nucleoids were similarly condensed following expression of toxic aberrant secretory proteins, suggesting that transertion of inner membrane proteins, a process that expands nucleoids into an open conformation that promotes growth and gene expression, was disrupted in these cells. The 48-residue ARF48 protein would be well-suited for structural studies to further investigate the toxic mechanism of aberrant proteins.

## Introduction

Aminoglycoside antibiotics such as kanamycin (Kan) bind to 16S rRNA in the A site of the 30S ribosomal subunit, interfering with the selection of cognate tRNAs [1]. Aberrant proteins are produced as a result, including proteins truncated by premature termination or with sequences garbled by frame-shift events or misincorporation of amino acids [2]. By contrast, protein synthesis is completely halted by non-aminoglycoside antibiotics like tetracycline (Tc) [3, 4] and chloramphenicol (Cm) [5]. Importantly, the growth-inhibiting effects of Tc and Cm are reversible, indicating that cells can tolerate absence of new protein synthesis for extended periods, whereas aminoglycosides are bactericidal [6]. Cells survive exposure to aminoglycosides if simultaneously treated with translation halting antibiotics like Cm, therefore indicating that the aberrant proteins synthesized in aminoglycoside treated cells are the ultimate cause of the bactericidal effect [6, 7]. Since each affected ribosome likely synthesizes a unique set of aberrant proteins, it has not been possible to directly study the sequence and structural determinants of aberrant protein toxic potential. Earlier work suggested that the diverse population of aberrant proteins may act in concert to disrupt cell membrane integrity, based on the finding that membrane permeability to ions increased following exposure to aminoglycoside antibiotics [8]. This led to the suggestion that aberrant proteins might be incorporated into the membrane, forming ion-conducting channels [7]. However, these putative channels were not isolated or directly characterized.

For aberrant proteins to have toxic effects they must escape detection and destruction by the cell’s protein quality control (QC) machinery. The heat shock (HS) response is the major protein QC system in bacteria and consists of a set of chaperones and proteases, together known as HS proteins, that function to rid cells of excess unfolded proteins and proteins that are irreparably damaged by exposure to extreme environmental conditions, such as elevated temperature [9]. HS gene expression is closely regulated to enable cells to adapt to changes in the burden of unfolded protein. The HS response is positively regulated by the transcription factor σ32 [10] and negatively regulated by the molecular chaperone DnaK, which binds σ32, promoting its degradation by the FtsH protease [11, 12].

Here we report that σ32 levels were increased about 10-fold in cells exposed to the aminoglycoside kanamycin (Kan), indicating that at least some aberrant proteins produced in these cells were recognized as substrates by DnaK. To gain insights into whether aberrant proteins vary significantly in their recognition as substrates by the QC machinery and whether this might be an important factor in determining their toxic potential, we studied aberrant proteins that were expressed in *E. coli* from cloned genes. Our results indicated that aberrant protein toxicity and recognition by the QC machinery are not mutually exclusive properties, suggesting that the QC machinery capacity to process aberrant proteins can be exceeded when aberrant proteins are produced at high rates. Our results further suggest that the toxic potential of aberrant proteins varies with their ability to disrupt the coupled transcription and co-translational insertion (transertion) of inner membrane (IM) proteins into the cell membrane, which is thought to stretch the nucleoid into an expanded conformation in growing cells [13-15].

## Materials and methods

### Protein expression constructs

The toxic ARF48 protein is encoded by an open +1 alternate reading frame (ARF) in an *Arabidopsis thaliana* cDNA for a Golgi-localized protein (At3G62370) involved in cell wall synthesis [16], which was obtained from the *Arabidopsis* Biological Resource Center at Ohio State University (clone U10771). A fragment of cDNA coding for the plant protein ectodomain (residues 18-325) was subcloned between the SacII and PstI sites in pASK-IBA3 (www.iba-lifesciences.com) and had an acutely toxic effect when expressed in *E. coli*. Deletion studies implicated ARF48 as the toxic factor, which was confirmed by subcloning the ARF48 coding sequence behind the TetA promoter in pASK-IBA3. Expression constructs for the ARF-NR and ARF-DA variants were created by PCR on circular ARF48 expression plasmid template DNA, using divergent primers that annealed to sites flanking the KL decapeptide coding sequence and had 5′ extensions that encoded the NR or DA heptapeptide substitutions, followed by self-ligation of the amplimers.

The coding sequence for *E. coli* alkaline phosphatase (AP) was amplified from genomic DNA and cloned behind the TetA promoter in pASK-IBA3 as previously described [17]. To create expression constructs for truncated AP variants, tandem stop codons were introduced in the AP reading frame between pre-AP codons 31-32, 149-150, 241-242, 337-338, or 429-430, using a 2-step method. In the first step, circular AP expression plasmid template DNA was amplified by PCR using divergent primer sets that annealed at each insertion point and had 5′ extensions specifying restriction sites for BamHI and XhoI. In the second step, the linear amplimers were doubly digested with BamHI and XhoI and were then ligated to a double-stranded oligonucleotide adapter that introduced tandem stop codons into the AP open reading frame. PCR with divergent-facing primers also was used to delete the coding sequence for pre-AP residues 10-14, to introduce the Δhc mutation into each truncated AP expression construct. All enzymes used for DNA manipulations were obtained from New England Biolabs, and the structure of all expression constructs was confirmed by DNA sequencing.

### Growth analysis

All growth studies were performed in LB broth containing antibiotic, using *E. coli* BL21 as the host strain except where noted. Three independent colony isolates of each construct were analyzed in parallel, and representative results are shown. Frozen stocks of cell strains were first grown overnight at 30°C, without shaking, in 2 ml of LB containing antibiotic. The starter cultures were then diluted 1:100 in 3.5 ml of LB containing antibiotic, in plastic 15×90 mm tubes with loose-fitting caps, and the tubes were then tilted about 15-20° from vertical and incubated at 37°C with shaking (280 rpm). Cell growth was monitored at the indicated times by measuring the optical density of culture samples (diluted 1:10 in saline) at 600 nm, using a Perkin Elmer Lambda 14 UV/VIS spectrophotometer and semi-micro glass cuvettes (1 ml sample volume). Expression of cloned genes from the Tet promoter in pASK-IBA3 was induced in early log phase cultures (OD_600_ ∼ 0.35) by addition of 0.2 µg/ml anhydrotetracycline (aTc) to the culture media. To measure cell viability, culture samples were serially diluted in LB broth, mixed with molten top agar and plated on LB agar containing antibiotic. The number of colonies was then counted after overnight incubation of plates at 37°C.

### Protein gel electrophoresis

Culture samples were centrifuged, and cell pellets were resuspended directly in 2X Laemmli sample buffer at a concentration of 5 OD_600_ units per ml. To reduce viscosity, samples were sonicated using a cuphorn sonicator or passed through a 27 gauge needle prior to loading on SDS-polyacrylamide gels. Proteins were visualized either by staining gels directly with Coomassie Blue or after semi-dry electroblotting to immobilon membranes for Western blot analysis. Anti-σ32 monoclonal antibodies were purchased from BioLegend, San Diego, CA. Rabbit anti-DnaK and DnaJ serum was kindly provided by B. Bukau. Anti-GroEL antibodies were purchased from Sigma-Aldrich.

### Peptide binding assay

Peptides were synthesized commercially (Peptide 2.0, Inc., Chantilly, VA) with fluorescein-modified N-terminal extensions (FITC-TEKA-) and amidated C-terminal extensions (-KTEQ-NH_2_). The concentrations of peptide solutions were determined by measuring fluorescein absorbance at 485 nm and applying the fluorescein extinction coefficient (50,358/M-cm). 6His-DnaK was purified from the ASKA expression clone [18] by chromatography on Ni-NTA agarose. The concentration of DnaK solutions was determined by BCA assay. 20 nM peptide was incubated with varying concentrations of DnaK in MST buffer (50 mM Tris pH 7.4, 150 mM NaCl, 10 mM MgCl_2_ supplemented with 0.05% Tween-20 and 0.05% bovine serum albumin) for 1 h at 20°C, before analysis by microscale thermophoresis [19], using a NanoTemper Technologies Monolith NT.115 instrument. Plots show the average ± SD for 3 independent trials for each peptide. Dissociation constants were determined using software provided with the instrument.

### RNA polymerase pull-down assay

An *E. coli* strain that produces RNA polymerase containing a 6-histidine-tagged β′ subunit [20] was obtained from M. Kashlev (NIH/NCI) and was transformed with the expression plasmids for ARF48 and the ARF-NR and ARF-DA variant proteins. Cells were grown to early log phase in 50 ml of LB broth containing ampicillin and were then treated with aTc to induce ARF polypeptide expression. At the indicated times 12.5 ml samples of culture were taken and the cells were lysed essentially as described [20]. Briefly, cells were washed in 1 ml of buffer 1 (10 mM Tris pH 8 / 100 mM NaCl / 1 mM β-mercaptoethanol / 5% glycerol), disrupted by sonicating 2x 15s with a microprobe in 300 µl of buffer 1 and then centrifuged 5min in a microcentrifuge. Protein concentration in supernatants was determined by BCA assay, and 100 µg samples of each lysate were then adjusted to 750 mg/ml by dilution with buffer 1. Samples were then adjusted to 0.05% Tween20 and 1 mM imidazole and incubated for 1 h on ice with 20 µl of Super Ni-NTA beads (www.proteinark.com), with frequent mixing. Beads were pelleted by centrifugation for 1 min at 830x g in a swinging bucket rotor, the unbound fraction (supernatant) was saved, and the beads were then washed twice in 100 µl buffer 1 containing 10 mM imidazole, then once in 100 µl buffer 1 containing 40 mM glycine. Bound proteins were then eluted in 50 µl of buffer 1 containing 150 mM imidazole.

Proteins in 15 µl samples of the pull-down and flow-through fractions were then run on 7% polyacrylamide gels and either stained with Coomassie Blue or transferred to PVDF membranes for Western blot analysis. Membranes were blocked for > 1h in PBS-T containing 5% BSA and then probed for 1 h with antibodies diluted in PBS-T containing 0.5% BSA. Mouse monoclonal antibodies against *E. coli* σ70 and σ32 were purchased from BioLegend (SanDiego, CA). Rabbit anti-GroEL IgG was purchased from Sigma-Aldrich. Rabbit sera against *E. coli* DnaK and DnaJ proteins was kindly provided by B. Bukau.

### Microscopy

Culture samples (0.3 ml) collected at the indicated times were centrifuged and the cell pellets were resuspended in 20-100 µl of Tris-buffered saline pH 8 (TBS) and held on ice. Cell suspensions were then stained with 1 µl of Hoechst 33342 (10 mg/ml stock solution was first diluted 1:100 in TBS). After incubation for 10 min on ice, 2 µl samples of cell suspension were mounted on coverslides using agarose blocks as described [21]. Cells were then examined by epi-fluorescence microscopy, using a Zeiss Axiophot and a 63X oil immersion lens. Exposure times for 2-channel images (both transmitted and reflected light) were set automatically using on-board Zen 2011 software. In some experiments as noted, exposure times for reflected light images were fixed at 1 s, for comparison of relative fluorescence intensity.

## Results

### Aberrant proteins produced in aminoglycoside-treated *E. coli* are substrates for DnaK

The molecular chaperone DnaK plays a key role in protein quality control (QC) by binding reversibly to exposed hydrophobic regions in unfolded proteins, blocking non-specific interactions that could disrupt protein folding [22]. DnaK also binds the heat shock (HS) transcription factor σ32 [11], but in contrast to the positive effect on most substrate proteins, interaction with DnaK promotes degradation of σ32 by proteolysis [23]. DnaK-dependent proteolysis of σ32 is competitively inhibited during periods of increased synthesis of DnaK substrate proteins, resulting in transient accumulation of σ32 protein and activation of the HS response [24]. We therefore monitored σ32 concentration to determine whether aberrant proteins synthesized in aminoglycoside-treated cells were recognized as substrates by DnaK.

Early log phase *E. coli* cells were exposed to the aminoglycoside kanamycin (Kan), and to the non-aminoglycoside antibiotics tetracycline (Tc) and chloramphenicol (Cm). Cell division was halted soon after exposure to all three antibiotics over a range of concentrations, as shown by leveling of the growth curves (Fig 1, panels A-C). σ32 levels in cells after 20, 60 or 120 min of antibiotic exposure were then determined by Western blotting. σ32 accumulated to high relative concentration in cells exposed to Kan, even at a low dose (3 µg/ml) that slowed but did not completely halt cell division, whereas σ32 was not detected in cells exposed to Cm or to any but the highest concentration (143 µg/ml) of Tc (Fig 1D; only the 60 min time point is shown). Quantification of the blotting data (Fig S1) showed that σ32 levels in Kan-treated cells increased by up to 12-fold in comparison to untreated control cells. Importantly, in contrast to the transient increase in σ32 concentration that occurs during an effective HS response, σ32 concentration remained permanently elevated in cells exposed to Kan (Fig 1E). These results indicated that aberrant proteins synthesized in Kan-treated cells can be recognized as substrates by DnaK, raising the question whether toxic species therefore might escape recognition or degradation by QC factors. The permanent increase in σ32 concentration also suggested that some step downstream of σ32 stabilization required for increased production of pro-survival HS proteins was blocked in these cells.

**Fig 1.**
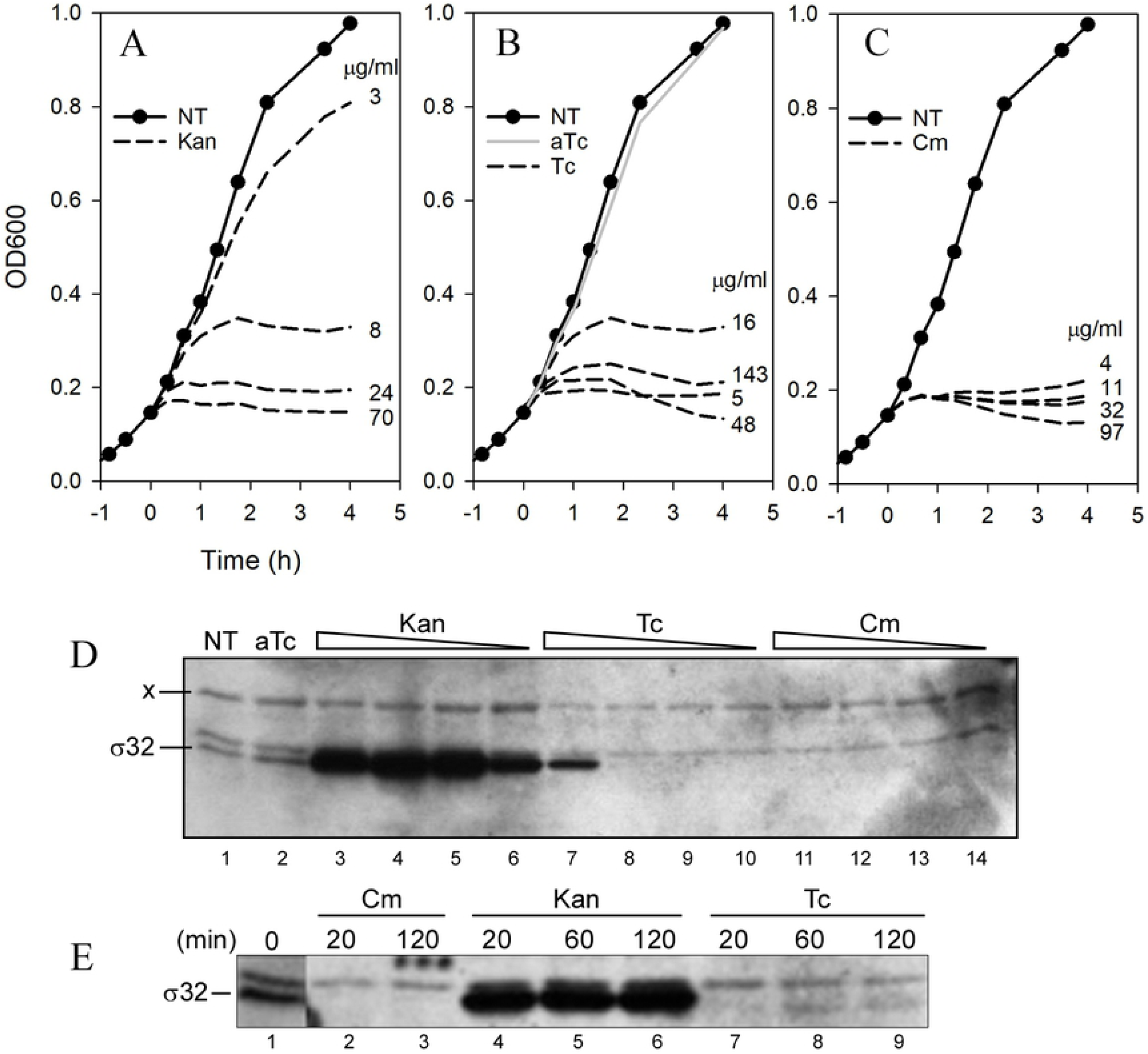
Effect of ribosome-targeting antibiotics on cell growth and σ32 stability. A-C, early log phase cultures of *E. coli* BL21 cells were exposed to the indicated doses of Kan, Tc or Cm (panels A, B and C, respectively), and cell division was then monitored by measuring culture optical density at 600 nm (OD600). D, an equal number of cells harvested from each culture at 60 min after exposure to antibiotics was analyzed by Western blotting for levels of the σ32 transcription factor. x, a cross-reacting protein that served as a loading control; NT, lysate from untreated control culture; aTc, lysate from culture exposed to 0.2 µg/ml anhydrotetracycline; triangles above lanes, relative dose of antibiotics as shown in panels A-C. E, Western blot analysis of σ32 concentration in cells following exposure to the indicated antibiotics for 20, 60, or 120 min as indicated. Cell extract from a non-induced culture was loaded in lane 1.

### A gene-encoded toxic aberrant protein

Each affected ribosome in aminoglycoside-treated cells likely synthesizes a unique set of aberrant proteins, and it is therefore difficult to isolate and characterize individual toxic species from these cells. To circumvent this problem, we studied aberrant proteins that were expressed in *E. coli* from cloned genes and mimicked the bactericidal effect of aminoglycoside antibiotics. ARF48 is a 48-residue protein encoded by an alternate reading frame in an *Arabidopsis thaliana* cDNA clone and is translated in *E. coli* from an internal Shine-Dalgarno-like sequence (Fig 2A-C). Cell division was halted immediately following ARF48 expression from a multi-copy plasmid (Fig 2D) and cell viability decreased about 10,000-fold by 2 h after induction (Fig 2E). ARF48 expression therefore mimics the bactericidal effect of aminoglycoside antibiotics [6]. By contrast, cells remained viable for several hours following growth arrest by 30 µg/ml of the bacteriostatic antibiotic Cm (Fig 2E). At higher concentration, Cm was increasingly toxic (Fig 2E).

**Fig 2.**
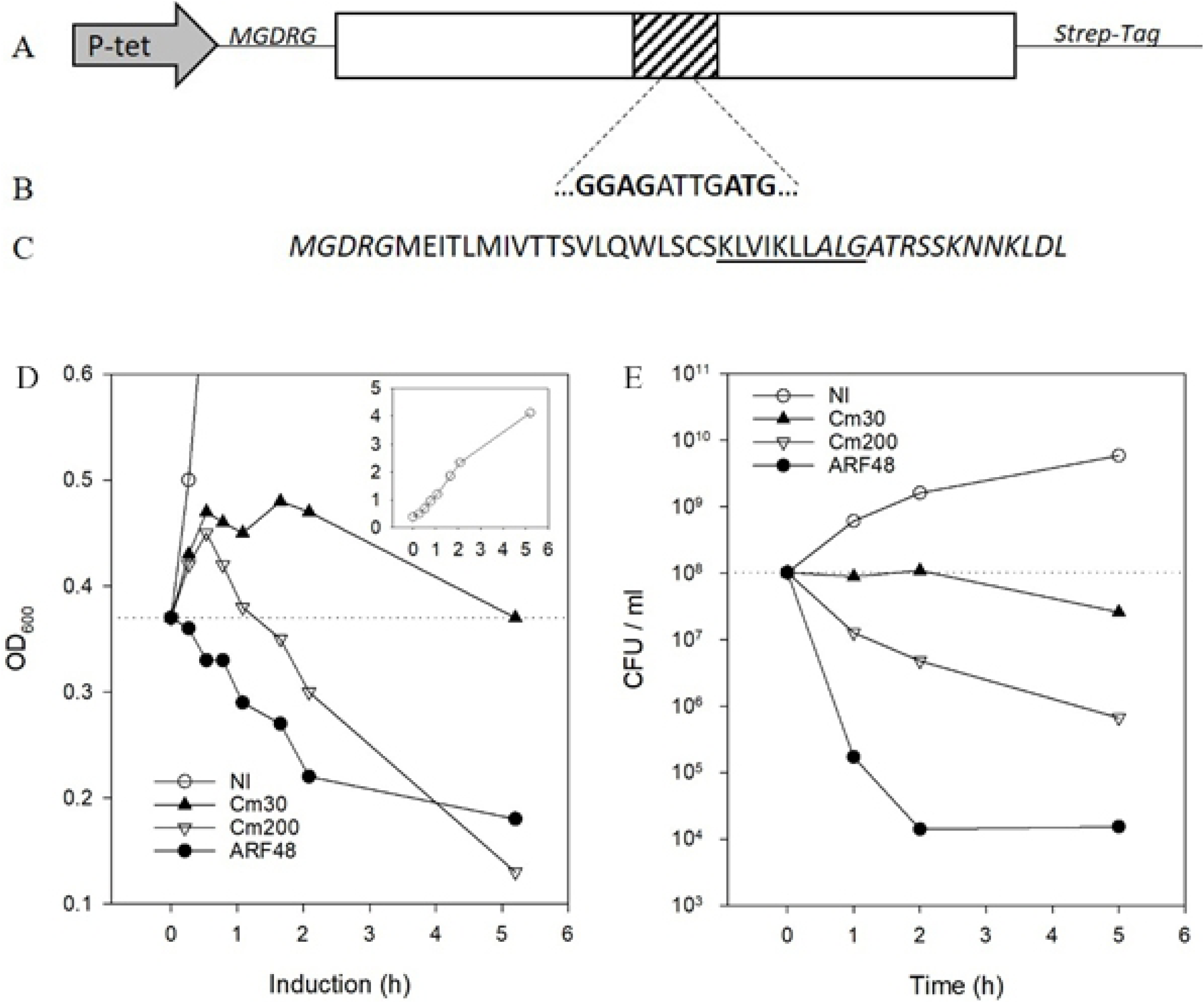
An alternate reading frame-encoded polypeptide is toxic when expressed in *E. coli*. A, an *Arabidopsis thaliana* cDNA (Genbank accession At3G62370; open rectangle) that was acutely toxic when expressed in *E. coli* from the tetracycline-inducible promoter (P-tet arrow) in pASK-IBA3. The plant protein coding sequence was fused in-frame to vector sequences coding for an N-terminal pentapeptide (MGDRG) and a C-terminal StrepTag. Deletion studies showed that an internal region of cDNA (shaded box, not drawn to scale) was sufficient for the toxic effect. B, a Shine-Dalgarno-like sequence [25] contained within the toxic region of cDNA. The ATG initiation codon for this putative translation start site is positioned in the +1 alternate reading frame (ARF) relative to the cDNA-encoded plant protein. C, the 48-residue protein (ARF48) produced by translation of the +1 ARF coding sequence from the start site in pASK-IBA3. A 10-residue sequence (KL decapeptide) that was tightly bound by DnaK in vitro is underlined. Vector-encoded residues are shown in italic type. D, the effect of ARF48 expression or exposure to 30 or 200 µg/ml Cm on growth of *E. coli* BL21 cells. NI, a non-induced culture grown in parallel, for comparison; inset, growth of the NI control culture. E, the effect of ARF48 expression or exposure to Cm on cell viability. Samples of culture taken at 0, 1, 2 and 5 h after treatment were serially diluted and plated on LB agar without inducer or Cm. Colonies were counted after incubating the plates overnight at 37°C. The results shown are representative of 3 independent experiments.

### The ARF48 toxic effect is gene dosage dependent

Proteins expressed from genes cloned in multi-copy plasmid vectors, such as the ColE1-based plasmid used for ARF48 expression, often are produced at high rates, leading to rapid accumulation of the proteins to high concentration [26]. To determine whether overexpression caused the ARF48 acutely toxic effect, we reduced the expression plasmid copy number by transforming cells with a second ColE1-based plasmid that conferred resistance to a different antibiotic [27]. The toxic effect indeed was attenuated when ARF48 was induced in cells that contained the co-resident ColE1 plasmid (Fig 3A). DNA gel electrophoresis confirmed that cells contained approximately equal amounts of each co-resident plasmid (Fig 3B).

**Fig 3.**
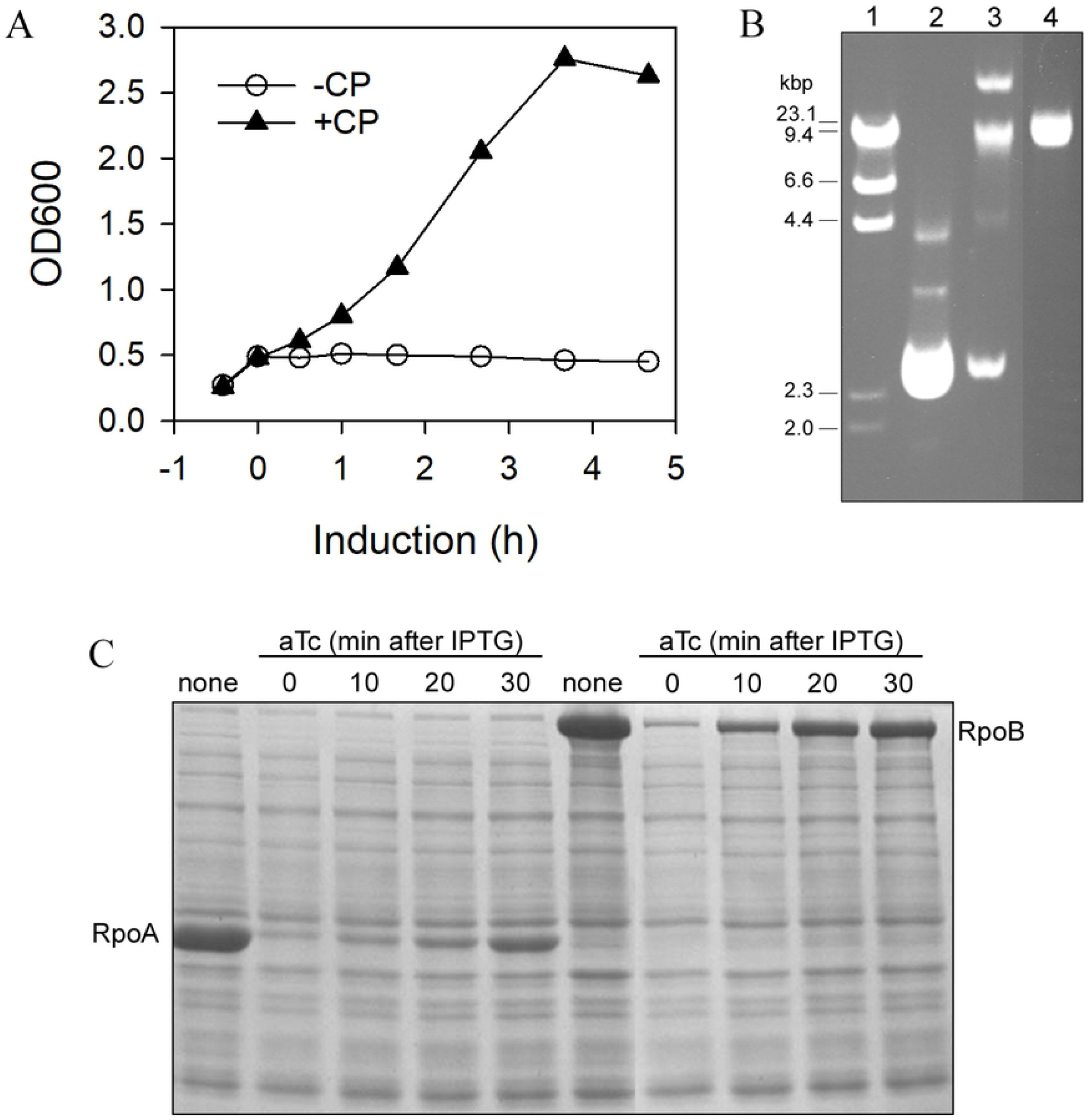
Effect of co-resident plasmids on the ARF48 toxic effect. A, cell growth following ARF48 induction (at T=0) in *E. coli* BL21(DE3) cells transformed by pARF48 alone (-CP, open circles) or in cells containing the co-resident plasmid pVS10(Sp-r), which confers resistance to spectinomycin (+CP, solid triangles) and is from the same plasmid incompatibility group as pARF48. B, agarose gel electrophoresis of plasmid DNA recovered from cells containing the co-resident plasmids pARF48 and pVS10(Sp-r) was loaded in lane 3. Purified samples of pARF48 and pVS10(Sp-r) were loaded in lanes 2 and 4, respectively. Phage lambda DNA digested with HindIII was loaded in lane 1, and the sizes of restriction fragments is indicated at left. C, *E. coli* BL21DE3 cells co-transformed by pARF48 and p15A-based expression plasmids for either RpoA or RpoB (different plasmid incompatibility groups) were grown to early log phase and then treated with IPTG (time 0) to induce expression of RpoA or RpoB. After incubation for 0, 10, 20 or 30 min, samples of each culture were removed and treated with aTc to induce ARF48 expression. After further incubation for 2 h, equal numbers of cells from each culture were lysed and the proteins were analyzed by SDS-polyacrylamide gel electrophoresis and Coomassie blue staining. Samples from control cultures not treated with aTc were loaded in lanes 1 and 6. The positions of RpoA and RpoB proteins in the gel are indicated. The corresponding growth curves for this experiment are shown in Fig S2.

In contrast to these results, the toxic effect was not attenuated when ARF48 was produced in cells containing co-resident plasmids from a different plasmid incompatibility group (p15A replicon; Fig S2). This was expect since the two plasmid replicons (ColE1 and p15A) in this case are regulated independently, and the coresident p15A plasmid therefore should not affect the ColE1-based ARF48 expression plasmid copy number [27]. Interestingly, in addition to halting cell growth, ARF48 induction also halted synthesis of the proteins expressed from genes cloned in the co-resident p15A plasmids (Fig 4C). Synthesis of LacZ (β-galactosidase), which is encoded by a single-copy gene localized in the chromosome, also was blocked following ARF48 expression (Table 1). Protein synthesis therefore appears to be globally halted following ARF48 overexpression.

**Table 1.** ARF48 blocks expression of β-galactosidase.

| Hours<br>after IPTG | +ARF48<sup>a</sup> | -ARF48<sup>b</sup> |
| --- | --- | --- |
| 1 | 3.0 ± 17% | 472 ± 22% |
| 2 | 13.8 ± 16% | 1137 ± 7% |

Cells were treated with aTc to induce ARF48 and were then treated with IPTG 5 min later to induce expression of β-galactosidase. After culturing for an additional 1 or 2 h, the amount of β-galactosidase enzyme activity (Miller units) in extracts of equal numbers of cells was determined.
<sup>a</sup> activity in cells expressing ARF48 (Miller units)
<sup>b</sup> activity in non-induced control cells

**Fig 4.**
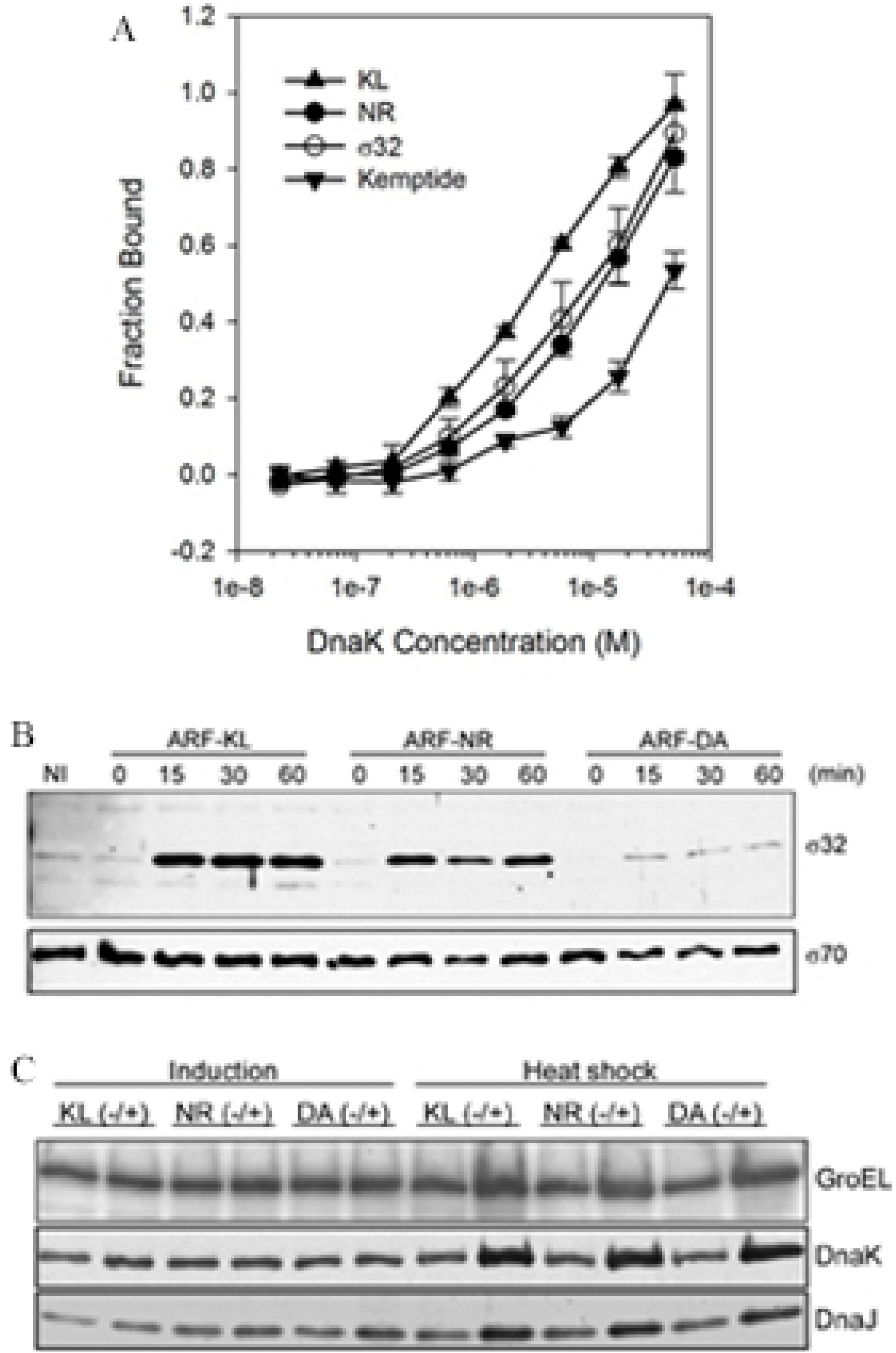
ARF48 is a substrate for DnaK in vitro and in vivo. A, equilibrium binding of fluorescein-labeled synthetic peptides (20 nM) to increasing concentrations of purified *E. coli* DnaK protein, determined by microscale thermophoresis [19]. All synthetic peptides were flanked by the same N- and C-terminal extensions (FITC-TEKA- and -KTEQ, respectively). KL, FITC-TEKAKLVIKLLALGKTEQ; NR, FITC-TEKANRLLLTGKTEQ; σ32, FITC-TEKAQRKLFFNLRKTEQ; Kemptide, FITC-TEKALRASLGKTEQ. Earlier studies showed that the NR and σ32 peptides were bound by DnaK in vitro [29, 31], and that Kemptide [32] was poorly recognized by comparison [33]. The results shown are representative of 3 independent experiments. D, the σ32 concentration in BL21 cells at 0, 15, 30 or 60 min after induction of ARF48 (ARF-KL) or the ARF-NR and ARF-DA variants, determined by Western blotting. NI, a non-induced control culture. Levels of the major sigma factor σ70 were determined in parallel (lower panel). The results shown are representative of 3 independent experiments. E, heat shock protein levels in cells before and after ARF protein induction (lanes 1-6) or heat shock at 42°C for 15 min (lanes 7-12).

### ARF48 is recognized as a substrate by DnaK

The foregoing results could be explained if the cell’s QC machinery recognizes ARF48 as a substrate and the machinery capacity to process aberrant proteins is exceeded when ARF48 is overexpressed. To test this model, we analyzed the ARF48 amino acid sequence for potential binding sites for DnaK, using the program *Limbo* [28]. *Limbo* predicted two partially overlapping binding sites for DnaK contained within ARF48 residues 25-34 (KLVIKLLALG; hereafter the KL decapeptide; underlined in Fig 2C). Purified DnaK indeed bound synthetic KL decapeptide tightly in vitro, with about 3-fold higher affinity than the canonical DnaK peptide substrate NRLLLTG [29, 30], as determined by microscale thermophoresis [19] (Fig 4A). Furthermore, the ARF48 toxic effect was attenuated or abrogated, respectively, when the KL decapeptide was substituted by NRLLLTG (ARF-NR) or by DAGAKAG (ARF-DA), a sequence predicted not to interact with DnaK (Fig S3A). The KL decapeptide therefore was recognized by DnaK in vitro and was required for the ARF48 toxic effect.

To determine whether ARF48 was recognized as a substrate by DnaK in vivo, we monitored σ32 concentration as described above. As in Kan-treated cells, the σ32 concentration increased following ARF48 induction and then remained at elevated concentration indefinitely, presumably until cell death (Fig 4B). σ32 concentration increased to a lesser degree following ARF-NR induction but did not increase following induction of ARF-DA (Fig 4B). ARF48 and ARF-NR therefore were recognized as substrates by DnaK in vivo. These proteins also were toxic when overexpressed in an *E. coli* mutant that lacks functional DnaK [34] however (Fig S3B), indicating that DnaK itself was not required for the toxic effect. These results support the model that the QC machinery capacity to process aberrant proteins is exceeded when ARF48 is overexpressed, and further suggest that the KL decapeptide mediates ARF48 interaction with some other factor(s) in addition to DnaK, which leads to cell death.

### ARF48 triggers an abortive HS response

Stabilization of σ32 typically indicates that HS gene expression has been activated [10], which normally would result in increased production of HS proteins that promote cell survival during periods of proteotoxic stress. However, cell viability decreased following ARF48 induction (Fig 2E) despite the increased concentration of σ32 in these cells, therefore indicating that the HS response was not effective. To further investigate this, we monitored the concentration of several HS proteins, including DnaK, DnaJ and GroEL, following ARF48 induction. Levels of these HS proteins essentially were unchanged following ARF48 induction but increased markedly when non-induced cultures were grown at 45°C for 10 min (Fig 4C). The HS response therefore functioned normally in these cells prior to ARF48 induction, suggesting that some step downstream of σ32 stabilization must be blocked following ARF48 induction.

To investigate whether the stabilized σ32 that accumulated following ARF48 expression could associate with RNA polymerase, we expressed ARF48 in an *E. coli* strain that produces RNA polymerase containing a 6-histidine-tagged β′ subunit [20]. Following ARF48 induction, RNA polymerase was purified from cell lysates by metal affinity chromatography, and the sigma factors associated with the enzyme were then analyzed by Western blotting. RNA polymerase isolated from cells following ARF48 induction indeed was associated with an increased amount of σ32 protein (E-σ32) in comparison to cells expressing ARF-NR or ARF-DA (Fig 5), indicating that the stabilized σ32 that accumulates following ARF48 expression is functionally intact. The failure of cells to increase production of HS proteins following ARF48 expression therefore must result from a block in either transcription of HS genes by E-σ32 or translation of HS protein mRNAs by cell ribosomes.

**Fig 5.**
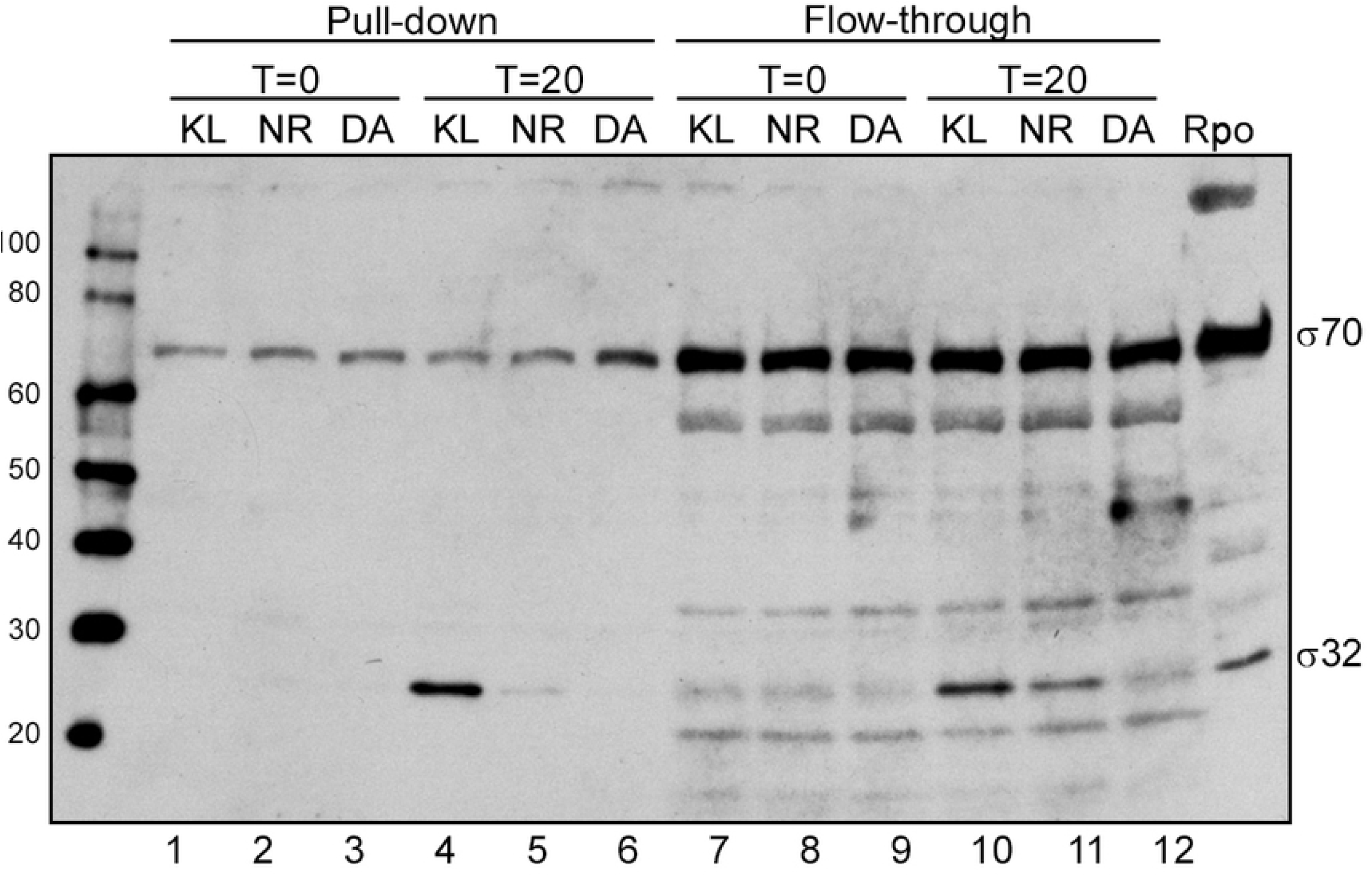
Excess σ32 that accumulates following ARF48 induction associates with RNA polymerase. ARF48 and the ARF-NR and ARF-DA variant polypeptides were expressed in an *E. coli* strain that produces RNA polymerase containing a 6-histidine-tagged β′ subunit [35]. Cell lysates taken immediately before induction (T=0) and 20 min after ARF protein induction (T=20) were incubated with Ni-NTA beads. Samples of the bound (Pull-down) and unbound (Flow-through) fractions were then electrophoresed in an SDS-polyacrylamide gel, transferred to an immobilon membrane, and then probed with antibodies specific for σ70 and σ32.

### ARF48 effect on nucleoid conformation

The initial decrease in culture OD600 following ARF48 induction (Fig 2D) suggested that some cell lysis might have occurred. However, the OD600 eventually stabilized (Fig 2D), and cell proteins were not detectably released into the culture medium (not shown). To further investigate the effect of ARF48 expression on cell membrane integrity, cells were incubated with the DNA-binding fluorescent dye Hoechst 33342 (H33342). Following ARF48 induction, cells accumulated approximately 8-times more H33342 than non-induced (NI) cells, based on the exposure times required to produce images of equal fluorescence intensity (Fig 6). Enhanced uptake of H33342 does not necessarily indicate the membrane is damaged however, because some bacteria can actively exclude H33342 by an efflux pumping mechanism [36]. Whether this efflux pumping mechanism also is present in *E. coli* has not yet been determined. Importantly, the microscopy also showed that nucleoids were radially condensed by 5-10 min after ARF48 induction, whereas the nucleoids in non-induced cells had a radially expanded conformation (Fig 6).

**Fig 6.**
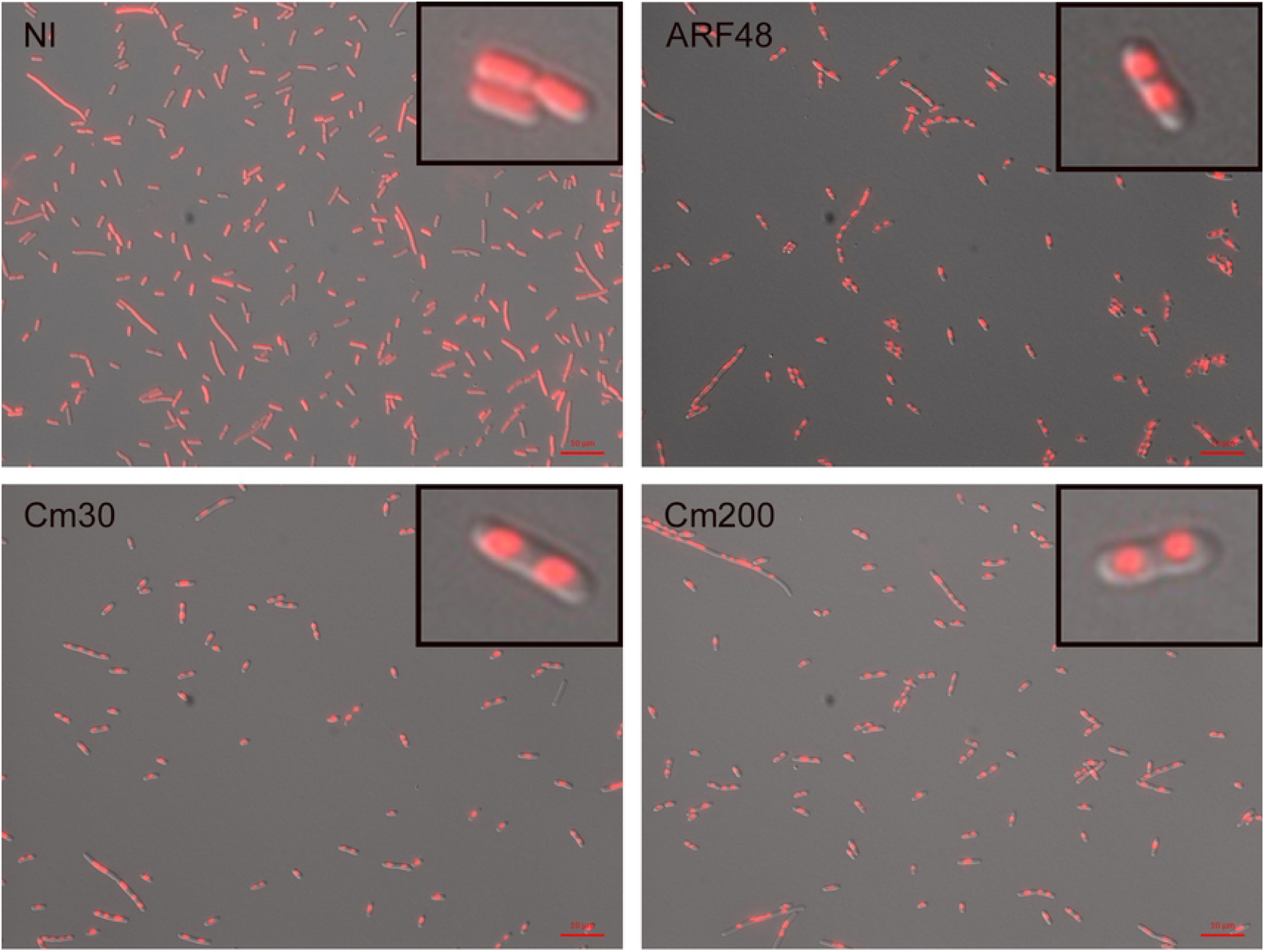
ARF48 induces radial condensation of nucleoids. An early log phase culture of BL21 cells transformed by pARF48 was divided and either left untreated (NI) or treated with anhydrotetracycline (aTc; 0.2 µg/ml) to induce ARF48 expression or with 30 or 200 µg/ml Cm. At 10 min after ARF48 induction or 60 min after treatment with Cm, cell samples were stained with H33342 and examined by fluorescence microscopy. Exposure times were automatically set by camera software and were 0.22 s for ARF48 and Cm samples and 1.7 s for the non-induced (NI) sample. Scale bar, 10 µm. The results shown are representative of at least 3 independent experiments.

Cm has long been known to induce radial condensation of nucleoids [37], which is thought to result from disruption of transertion, the coupled transcription and co-translational insertion of inner membrane (IM) proteins into the cell membrane [13, 15]. During transertion, the cognate IM protein genes are anchored to the cell membrane [38], thus stretching the nucleoid into an open or radially expanded conformation [14]. Consistent with earlier studies, we also found that nucleoids were radially condensed following exposure to Cm (Fig 6; Cm30 and Cm200), but the effect took longer to develop in comparison to ARF48 induction and was not uniformly apparent in all cells until 30-45 min after treatment. As occurred following ARF48 expression, uptake of H33342 also was enhanced following exposure to Cm. Since the effects of Cm (30 µg/ml) are reversible for several hours (Fig 2D), the enhanced uptake of H33342 in this case does not indicate that the cells are dead or that the membrane has been irreparably damaged.

### Sec pathway localization enhances aberrant protein toxic potential

IM proteins are inserted into the membrane through translocons, channels in the membrane formed by the SecYEG protein complex [39]. Earlier studies showed that cytoplasmic proteins such as LacZ (β-galactosidase) can be mis-localized into the protein secretion (Sec) pathway by fusing a signal peptide to the protein N-terminus, and that this can result in lethal jamming of translocons [40]. To determine the impact of translocon jamming on nucleoid conformation, we studied a LamB-LacZ fusion protein that previously was shown to jam translocons [41]. In contrast to the plasmid-borne ARF48 gene, the LamB-LacZ fusion protein gene is present in just one copy per chromosome. Following induction of LamB-LacZ expression, cell growth continued at a slow rate for about 3 h before finally halting (Fig 7A). Cell uptake of H33342 increased markedly after 1-2 h of induction, and by 4 h many nucleoids had a radially condensed conformation (Fig 7B).

**Fig 7.**
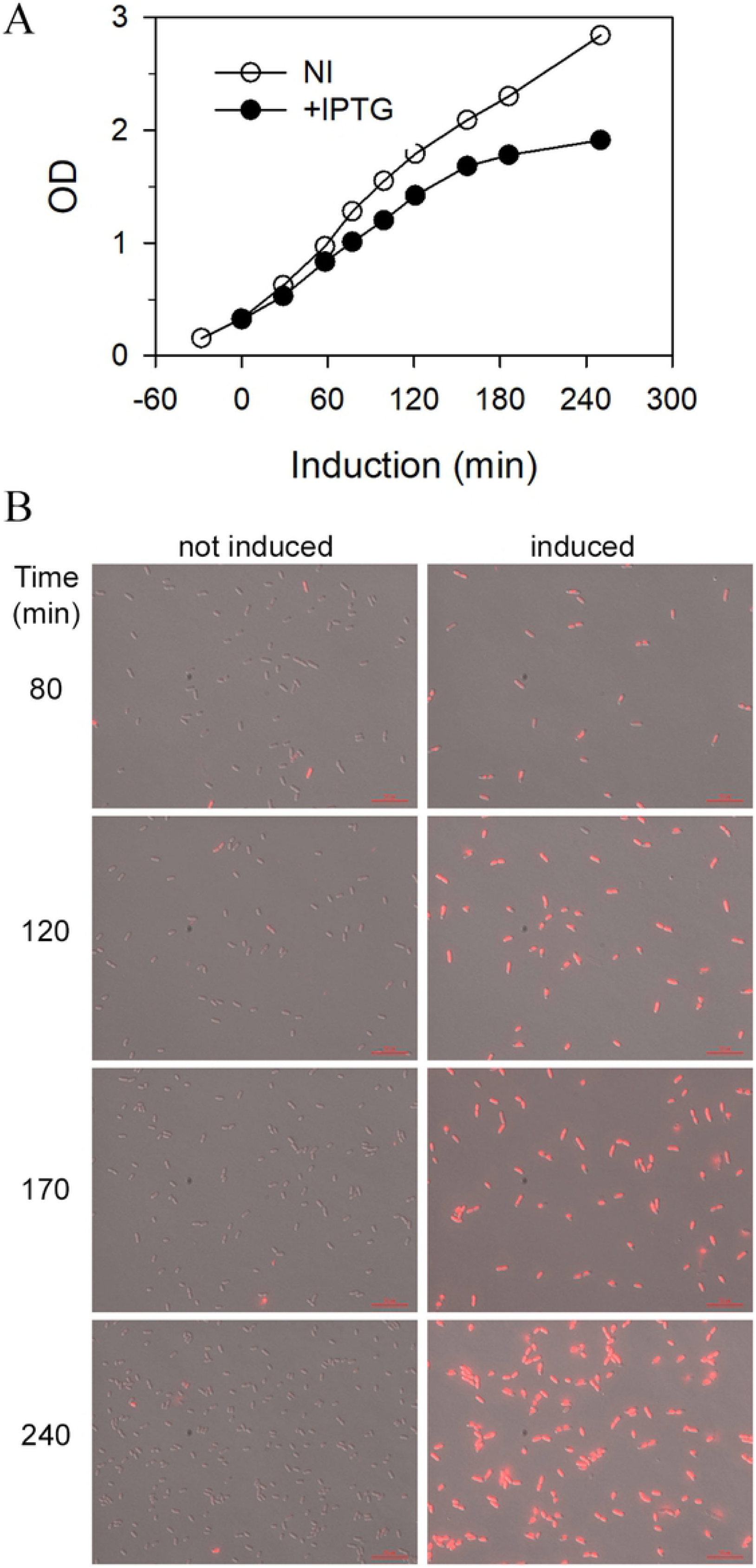
Toxic LamB-LacZ fusion protein increases cell uptake of H33342 and induces radial condensation of nucleoids. A, growth of *E. coli* strain JCM912 (kindly provided by T. Silhavy) following induction (by IPTG) of a toxic LamB-LacZ fusion protein expressed from a single-copy chromosomal gene (filled circles), in comparison to a non-induced control culture (open circles). B, fluorescence microscopy of cell samples from both cultures, taken at the indicated times after LamB-LacZ induction. The exposure time for all images was fixed at 1 s.

To further investigate the impact of Sec pathway localization on aberrant protein toxic potential and nucleoid conformation, we overexpressed truncated variants of the *E. coli* alkaline phosphatase (AP) protein. AP is a periplasmic enzyme that is synthesized as a precursor with an N-terminal signal peptide that is cleaved during or soon after translocation across the cell membrane [42]. Premature stop codons were introduced at different positions within the AP coding sequence, to mimic the protein truncating effect of aminoglycoside antibiotics. Cell growth was inhibited to varying degrees by all the truncated AP variants, but the smallest proteins, which were truncated after AP residues 149 or 241, had the most acutely toxic effect (Fig 8A). Enhanced accumulation of the larger variants (Fig 8C) suggests they may have self-aggregated before entering the Sec pathway. Deletion of 5 residues (LLPLL) from the signal peptide hydrophobic core (Δhc) attenuated the toxic effect of all truncated variants (Fig 8B), therefore indicating that Sec pathway localization is required for the toxic effect of these proteins. Nucleoids were rapidly condensed following expression of the acutely toxic variants (Fig 8E) and the effect was suppressed by the Δhc mutation (Fig 8D). These results support the conclusion that the toxic effect of aberrant proteins can result from disruption of transertion-mediated expansion of nucleoids into an open conformation required for cell growth [15].

**Fig 8.**
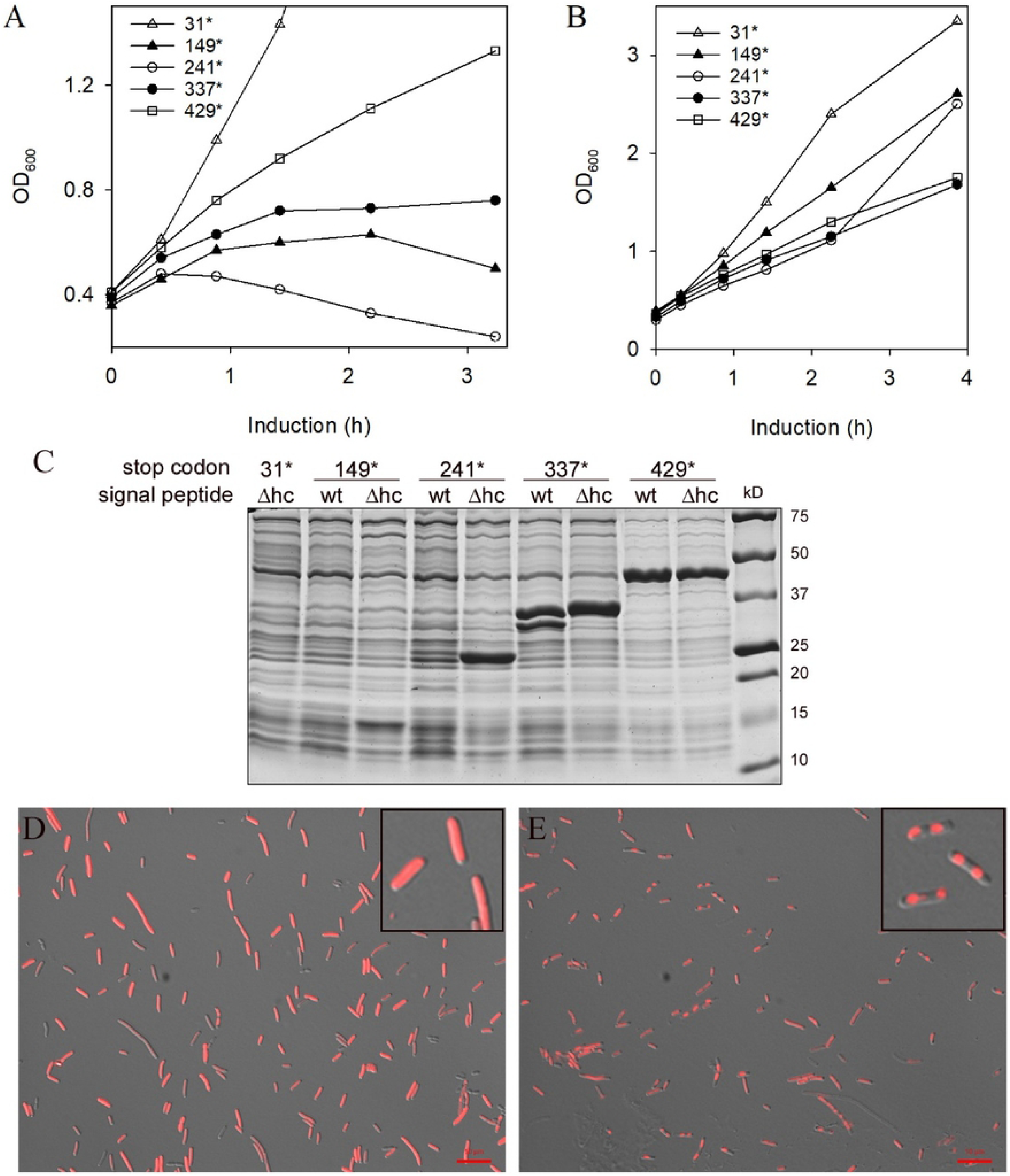
Truncated alkaline phosphatase (AP) variants are toxic when localized to the Sec pathway. A, growth of BL21 cells following induction of AP variants that were truncated after preAP codons 31, 149, 241, 337, or 429 but contain an intact N-terminal signal peptide. B, same experiment as in A except the signal peptide of each truncated AP variant was inactivated by deletion of 5 amino acid residues (LLPLL) from the hydrophobic core (Δhc). C, 2h after induction, cell samples from the indicated cultures were lysed in 2X Laemmli sample buffer and electrophoresed in an SDS-polyacrylamide gel. Proteins were then visualized by staining with Coomassie blue. D, nucleoid conformation 1 h after induction of AP241 containing a mutant signal peptide (Δhc). E, nucleoid conformation 1 h after induction of AP241 containing a wild-type signal peptide. Scale bar, 10 µm. Cells in insets are enlarged 3X. Exposure times were set automatically by microscope software and were 1.8 s (D) and 0.3 s (E).

## Discussion

Misfolded and aberrant proteins have toxic potential [6, 43], but the underlying mechanisms have proven difficult to elucidate. Here we show that the HS transcription factor σ32 concentration increases following treatment of *E. coli* with the aminoglycoside Kan, indicating that at least some of the diverse aberrant proteins synthesized in these cells are recognized as substrates by the molecular chaperone DnaK [24]. Stabilization of σ32 indicates that HS gene expression has been activated, which typically results in increased synthesis of HS proteins that restore protein homeostasis and reset σ32 concentration back to basal levels. σ32 concentration was permanently elevated following exposure to Kan however, suggesting that the HS response in these cells was blocked at some stage downstream of σ32 stabilization.

To investigate the abortive HS response in more detail without possible complicating effects of Kan or other antibiotics, we studied aberrant proteins that mimicked the bactericidal effect of aminoglycosides when expressed in *E. coli* from cloned genes. ARF48 proved to be an excellent model for this study, since it not only was bactericidal but also caused σ32 concentration to remain permanently elevated, as occurred following exposure to Kan. Knowledge of the ARF48 amino acid sequence and access to the coding sequence enabled the discovery of a 10-residue region, termed the KL decapeptide, that was required both for the toxic effect and for recognition by DnaK. This important finding indicates that toxic activity and recognition as a QC substrate are not mutually exclusive properties of aberrant proteins. This apparent paradox was resolved by the finding that the ARF48 toxic effect was proportional to gene dosage. Together these results suggest that QC capacity can be exceeded when aberrant proteins are produced at high rates, as can occur when aberrant proteins are overexpressed from genes cloned on multi-copy plasmid vectors or when aminoglycoside antibiotics are used above the minimum inhibitory concentration.

Our results also provided insights into the toxic mechanism of excess aberrant proteins that accumulate when QC capacity is saturated. As occurred following exposure of cells to Kan, σ32 concentration increased about 10-fold over basal levels following ARF48 induction and remained at this elevated level indefinitely. As noted above, this atypical, prolonged stabilization of σ32 suggests that later stages of the HS response downstream of σ32 stabilization were not properly executed. In support of this conclusion, we found that levels of several HS proteins did not increase following ARF48 induction. This block was not specific for HS proteins, because synthesis of other proteins unrelated to the HS response also was suppressed. Given the similarity to other effects of ribosome-targeting antibiotics, it is possible although unlikely that the global block in protein synthesis could result from direct inhibition of ribosomes by ARF48, in an antibiotic-like manner. In preliminary studies, we determined that the KL decapeptide is efficiently translated when inserted as a guest peptide at several different positions in alkaline phosphatase (AB and PF, unpublished observations), indicating that this sequence element is not retained in the ribosomal exit tunnel as can occur during translation of the *E. coli* SecM protein [44, 45].

Radial condensation of nucleoids is a more likely mechanism for the general block in protein synthesis that occurs following ARF48 expression. Nucleoids in growing cells have an expanded conformation thought to result from coupled transcription and co-translational insertion (transertion) of inner membrane (IM) proteins into the cell membrane [13-15]. Based on modeling studies [15], it was proposed that the expanded nucleoid conformation is necessary for recycling of the 30S and 50S ribosomal subunits back into the genome for coupling of transcription and translation. According to this model, the rapid condensation of nucleoids that occurs following ARF48 induction would block recycling of ribosomal subunits, thus globally inhibiting gene expression.

Consistent with the model that transertion mediates expansion of nucleoids in growing cells, nucleoids are radially condensed when transcription or translation are inhibited by antibiotics [13, 15]. Since translocons serve as portals for insertion of IM proteins into the cell membrane [39] they also should be essential for transertion-mediated nucleoid expansion, but a role for translocons in nucleoid conformation has not been reported. Here we showed that nucleoids were radially condensed following expression of a LamB-LacZ fusion protein that previously was shown to jam translocons [46]. Nucleoids were similarly condensed following overexpression of truncated variants of the secretory protein alkaline phosphatase, and the effect was dependent on signal peptide function. These results are consistent with the model that transertion mediates nucleoid expansion and provide the first reported evidence that translocons play an essential role in this process.

The radial condensation of nucleoids following ARF48 induction suggests that ARF48 also might be localized to the Sec pathway. Unlike AP and LamB-LacZ, ARF48 does not have an N-terminal signal peptide. However, our results indicate that the KL decapeptide mediates ARF48 interaction with an unidentified factor, in addition to DnaK, that might target ARF48 to the membrane. Possible candidate binding partners for KL include the signal recognition particle (SRP) and the SecA translocase, both of which were recently shown to have non-canonical interactions with substrate proteins [47, 48]. Earlier studies also showed that signal peptides enhance but are not always required for protein export through the Sec pathway [49]. Further studies will be needed to elucidate the ARF48 toxic mechanism. The small size of this protein would be advantageous for structural studies, which could reveal targets for novel antimicrobial drugs.

## Acknowledgments

We are grateful to D. Figurski for modifying pVS10 [50] to carry a gene conferring resistance to spectinomycin in place of the ampicillin-resistance gene; to M. Gottesman, M. Kashlev and T. Silhavy for generously providing *E. coli* strains; to B. Bukau for providing anti-DnaK and anti-DnaJ serum; to G. Guo and A. Kim for assistance with microscale thermophoresis and fluorescence microscopy; and to D. Figurski, M. Gottesman, C. Anderson, and D. Weiss for helpful discussions. S. Pearson and P. Muhanji provided excellent technical assistance.

## Supporting information

**S1 Fig. Effect of Kan on σ32 stability**. The data shown in Fig 1D (fold increase in σ32 concentration after 60 min of exposure to varying doses of Kan) were quantified using the program Image J.

**S2 Fig. The ARF48 toxic effect is not suppressed in cells containing co-resident plasmids from a different incompatibility group**. Shown are growth curves corresponding to the experiment shown in Fig 3C. A, cell growth following ARF48 induction in cells containing a co-resident expression plasmid for RpoA (pACYC-pET-RpoA). Solid line with open circles, growth of the parental culture expressing RpoA alone; dotted lines with filled triangles, growth of parental culture samples after addition of aTc to induce ARF48 expression. The parental culture was treated with IPTG at time 0 to induce expression of RpoA. B, same experiment as A, except ARF48 was induced in cells containing a co-resident expression plasmid for RpoB (pACYC-pET-RpoB).

**S3 Fig. DnaK is not required for the ARF48 toxic effect**. A, growth of E. coli BL21 cells (which contain wild-type DnaK) following expression of ARF48 or the ARF-NR and ARF-DA variants. △, non-induced control; ▴, ARF48; ○, ARF-NR; ●, ARF-DA. B, same experiment as A, but proteins were expressed in a mutant *E. coli* strain that lacks the DnaK gene (*ΔdnaK*) [34]. The results shown are representative of 3 independent experiments.

## References

1. Carter AP, Clemons WM, Brodersen DE, Morgan-Warren RJ, Wimberly BT, Ramakrishnan V. Functional insights from the structure of the 30S ribosomal subunit and its interactions with antibiotics. Nature. 2000;407(6802):340-8. Epub 2000/10/03. doi: 10.1038/35030019. PubMed PMID: 11014183.

2. Aguirre Rivera J, Larsson J, Volkov IL, Seefeldt AC, Sanyal S, Johansson M. Real-time measurements of aminoglycoside effects on protein synthesis in live cells. Proc Natl Acad Sci U S A. 2021;118(9). Epub 2021/02/24. doi: 10.1073/pnas.2013315118. PubMed PMID: 33619089; PubMed Central PMCID: PMCPMC7936356.

3. Brodersen DE, Clemons WM, Jr., Carter AP, Morgan-Warren RJ, Wimberly BT, Ramakrishnan V. The structural basis for the action of the antibiotics tetracycline, pactamycin, and hygromycin B on the 30S ribosomal subunit. Cell. 2000;103(7):1143–54. doi: 10.1016/s0092-8674(00)00216-6. PubMed PMID: 11163189.

4. Pioletti M, Schlünzen F, Harms J, Zarivach R, Glühmann M, Avila H, et al. Crystal structures of complexes of the small ribosomal subunit with tetracycline, edeine and IF3. Embo j. 2001;20(8):1829–39. doi: 10.1093/emboj/20.8.1829. PubMed PMID: 11296217; PubMed Central PMCID: PMCPMC125237.

5. Bulkley D, Innis CA, Blaha G, Steitz TA. Revisiting the structures of several antibiotics bound to the bacterial ribosome. Proc Natl Acad Sci U S A. 2010;107(40):17158-63. Epub 2010/09/30. doi: 10.1073/pnas.1008685107. PubMed PMID: 20876130; PubMed Central PMCID: PMCPMC2951403.

6. Baquero F, Levin BR. Proximate and ultimate causes of the bactericidal action of antibiotics. Nat Rev Microbiol. 2021;19(2):123-32. Epub 2020/10/08. doi: 10.1038/s41579-020-00443-1. PubMed PMID: 33024310; PubMed Central PMCID: PMCPMC7537969.

7. Davis BD, Chen LL, Tai PC. Misread protein creates membrane channels: an essential step in the bactericidal action of aminoglycosides. Proc Natl Acad Sci U S A. 1986;83(16):6164–8. doi: 10.1073/pnas.83.16.6164. PubMed PMID: 2426712; PubMed Central PMCID: PMCPMC386460.

8. Davis BD. Mechanism of bactericidal action of aminoglycosides. Microbiol Rev. 1987;51(3):341-50. Epub 1987/09/01. PubMed PMID: 3312985; PubMed Central PMCID: PMCPMC373115.

9. Arsene F, Tomoyasu T, Bukau B. The heat shock response of Escherichia coli. Int J Food Microbiol. 2000;55(1-3):3-9. Epub 2000/05/03. PubMed PMID: 10791710.

10. Straus DB, Walter WA, Gross CA. The heat shock response of E. coli is regulated by changes in the concentration of sigma 32. Nature. 1987;329(6137):348-51. Epub 1987/09/24. doi: 10.1038/329348a0. PubMed PMID: 3306410.

11. Liberek K, Galitski TP, Zylicz M, Georgopoulos C. The DnaK chaperone modulates the heat shock response of Escherichia coli by binding to the sigma 32 transcription factor. Proc Natl Acad Sci U S A. 1992;89(8):3516–20. doi: 10.1073/pnas.89.8.3516. PubMed PMID: 1565647; PubMed Central PMCID: PMCPMC48899.

12. Tomoyasu T, Gamer J, Bukau B, Kanemori M, Mori H, Rutman AJ, et al. Escherichia coli FtsH is a membrane-bound, ATP-dependent protease which degrades the heat-shock transcription factor sigma 32. Embo j. 1995;14(11):2551-60. Epub 1995/06/01. PubMed PMID: 7781608; PubMed Central PMCID: PMCPMC398369.

13. Roggiani M, Goulian M. Chromosome-Membrane Interactions in Bacteria. Annu Rev Genet. 2015;49:115-29. Epub 2015/10/06. doi: 10.1146/annurev-genet-112414-054958. PubMed PMID: 26436460.

14. Woldringh CL. The role of co-transcriptional translation and protein translocation (transertion) in bacterial chromosome segregation. Mol Microbiol. 2002;45(1):17-29. Epub 2002/07/09. PubMed PMID: 12100545.

15. Bakshi S, Choi H, Weisshaar JC. The spatial biology of transcription and translation in rapidly growing Escherichia coli. Front Microbiol. 2015;6:636. Epub 2015/07/21. doi: 10.3389/fmicb.2015.00636. PubMed PMID: 26191045; PubMed Central PMCID: PMCPMC4488752.

16. Parsons HT, Christiansen K, Knierim B, Carroll A, Ito J, Batth TS, et al. Isolation and proteomic characterization of the Arabidopsis Golgi defines functional and novel components involved in plant cell wall biosynthesis. Plant Physiol. 2012;159(1):12-26. Epub 2012/03/21. doi: 10.1104/pp.111.193151. PubMed PMID: 22430844; PubMed Central PMCID: PMCPmc3375956.

17. Voss S, Skerra A. Mutagenesis of a flexible loop in streptavidin leads to higher affinity for the Strep-tag II peptide and improved performance in recombinant protein purification. Protein Eng. 1997;10(8):975-82. Epub 1997/08/01. PubMed PMID: 9415448.

18. Kitagawa M, Ara T, Arifuzzaman M, Ioka-Nakamichi T, Inamoto E, Toyonaga H, et al. Complete set of ORF clones of Escherichia coli ASKA library (a complete set of E. coli K-12 ORF archive): unique resources for biological research. DNA Res. 2005;12(5):291-9. Epub 2006/06/14. doi: 10.1093/dnares/dsi012. PubMed PMID: 16769691.

19. Wienken CJ, Baaske P, Rothbauer U, Braun D, Duhr S. Protein-binding assays in biological liquids using microscale thermophoresis. Nature Communications. 2010;1:100. doi: 10.1038/ncomms1093 https://www.nature.com/articles/ncomms1093#supplementary-information.

20. Kashlev M, Martin E, Polyakov A, Severinov K, Nikiforov V, Goldfarb A. Histidine-tagged RNA polymerase: dissection of the transcription cycle using immobilized enzyme. Gene. 1993;130(1):9-14. Epub 1993/08/16. doi: 10.1016/0378-1119(93)90340-9. PubMed PMID: 8344532.

21. Skinner SO, Sepulveda LA, Xu H, Golding I. Measuring mRNA copy number in individual Escherichia coli cells using single-molecule fluorescent in situ hybridization. Nat Protoc. 2013;8(6):1100-13. Epub 2013/05/18. doi: 10.1038/nprot.2013.066. PubMed PMID: 23680982; PubMed Central PMCID: PMCPMC4029592.

22. Bukau B, Horwich AL. The Hsp70 and Hsp60 chaperone machines. Cell. 1998;92(3):351-66. Epub 1998/02/26. PubMed PMID: 9476895.

23. Rodriguez F, Arsene-Ploetze F, Rist W, Rudiger S, Schneider-Mergener J, Mayer MP, et al. Molecular basis for regulation of the heat shock transcription factor sigma32 by the DnaK and DnaJ chaperones. Mol Cell. 2008;32(3):347-58. Epub 2008/11/11. doi: 10.1016/j.molcel.2008.09.016. PubMed PMID: 18995833.

24. Kanemori M, Mori H, Yura T. Induction of heat shock proteins by abnormal proteins results from stabilization and not increased synthesis of sigma 32 in Escherichia coli. J Bacteriol. 1994;176(18):5648-53. Epub 1994/09/01. PubMed PMID: 7916010; PubMed Central PMCID: PMCPMC196767.

25. Shine J, Dalgarno L. Determinant of cistron specificity in bacterial ribosomes. Nature. 1975;254(5495):34-8. Epub 1975/03/06. PubMed PMID: 803646.

26. Studier FW, Moffatt BA. Use of bacteriophage T7 RNA polymerase to direct selective high-level expression of cloned genes. J Mol Biol. 1986;189(1):113-30. Epub 1986/05/05. PubMed PMID: 3537305.

27. Novick RP. Plasmid incompatibility. Microbiol Rev. 1987;51(4):381–95. doi: 10.1128/mr.51.4.381-395.1987. PubMed PMID: 3325793; PubMed Central PMCID: PMCPMC373122.

28. Van Durme J, Maurer-Stroh S, Gallardo R, Wilkinson H, Rousseau F, Schymkowitz J. Accurate prediction of DnaK-peptide binding via homology modelling and experimental data. PLoS Comput Biol. 2009;5(8):e1000475. Epub 2009/08/22. doi: 10.1371/journal.pcbi.1000475. PubMed PMID: 19696878; PubMed Central PMCID: PMCPMC2717214.

29. Gragerov A, Zeng L, Zhao X, Burkholder W, Gottesman ME. Specificity of DnaK-peptide binding. J Mol Biol. 1994;235(3):848-54. Epub 1994/01/21. doi: 10.1006/jmbi.1994.1043. PubMed PMID: 8289323.

30. Zhu X, Zhao X, Burkholder WF, Gragerov A, Ogata CM, Gottesman ME, et al. Structural analysis of substrate binding by the molecular chaperone DnaK. Science. 1996;272(5268):1606-14. Epub 1996/06/14. PubMed PMID: 8658133.

31. McCarty JS, Rudiger S, Schonfeld HJ, Schneider-Mergener J, Nakahigashi K, Yura T, et al. Regulatory region C of the E. coli heat shock transcription factor, sigma32, constitutes a DnaK binding site and is conserved among eubacteria. J Mol Biol. 1996;256(5):829-37. Epub 1996/03/15. PubMed PMID: 8601834.

32. Maller JL, Kemp BE, Krebs EG. In vivo phosphorylation of a synthetic peptide substrate of cyclic AMP-dependent protein kinase. Proc Natl Acad Sci U S A. 1978;75(1):248-51. Epub 1978/01/01. PubMed PMID: 203933; PubMed Central PMCID: PMCPMC411223.

33. Rudiger S, Germeroth L, Schneider-Mergener J, Bukau B. Substrate specificity of the DnaK chaperone determined by screening cellulose-bound peptide libraries. Embo j. 1997;16(7):1501-7. Epub 1997/04/01. doi: 10.1093/emboj/16.7.1501. PubMed PMID: 9130695; PubMed Central PMCID: PMCPMC1169754.

34. Baba T, Ara T, Hasegawa M, Takai Y, Okumura Y, Baba M, et al. Construction of Escherichia coli K-12 in-frame, single-gene knockout mutants: the Keio collection. Mol Syst Biol. 2006;2:2006.0008. Epub 2006/06/02. doi: 10.1038/msb4100050. PubMed PMID: 16738554; PubMed Central PMCID: PMCPMC1681482.

35. Kashlev M, Nudler E, Severinov K, Borukhov S, Komissarova N, Goldfarb A. Histidine-tagged RNA polymerase of Escherichia coli and transcription in solid phase. Methods Enzymol. 1996;274:326-34. Epub 1996/01/01. doi: 10.1016/s0076-6879(96)74028-4. PubMed PMID: 8902816.

36. Venter H, Velamakanni S, Balakrishnan L, van Veen HW. On the energy-dependence of Hoechst 33342 transport by the ABC transporter LmrA. Biochem Pharmacol. 2008;75(4):866-74. Epub 20071026. doi: 10.1016/j.bcp.2007.10.022. PubMed PMID: 18061142.

37. Morgan C, Rosenkranz HS, Carr HS, Rose HM. Electron microscopy of chloramphenicol-treated Escherichia coli. J Bacteriol. 1967;93(6):1987-2002. Epub 1967/06/01. PubMed PMID: 5337775; PubMed Central PMCID: PMCPMC276719.

38. Lynch AS, Wang JC. Anchoring of DNA to the bacterial cytoplasmic membrane through cotranscriptional synthesis of polypeptides encoding membrane proteins or proteins for export: a mechanism of plasmid hypernegative supercoiling in mutants deficient in DNA topoisomerase I. J Bacteriol. 1993;175(6):1645-55. Epub 1993/03/01. doi: 10.1128/jb.175.6.1645-1655.1993. PubMed PMID: 8383663; PubMed Central PMCID: PMCPMC203958.

39. Beckwith J. The Sec-dependent pathway. Res Microbiol. 2013;164(6):497-504. Epub 2013/03/30. doi: 10.1016/j.resmic.2013.03.007. PubMed PMID: 23538404; PubMed Central PMCID: PMCPMC3706482.

40. Bassford PJ, Jr., Silhavy TJ, Beckwith JR. Use of gene fusion to study secretion of maltose-binding protein into Escherichia coli periplasm. J Bacteriol. 1979;139(1):19-31. Epub 1979/07/01. PubMed PMID: 110778; PubMed Central PMCID: PMCPMC216822.

41. Dwyer RS, Malinverni JC, Boyd D, Beckwith J, Silhavy TJ. Folding LacZ in the periplasm of Escherichia coli. J Bacteriol. 2014;196(18):3343-50. Epub 2014/07/09. doi: 10.1128/jb.01843-14. PubMed PMID: 25002543; PubMed Central PMCID: PMCPMC4135689.

42. Josefsson LG, Randall LL. Different exported proteins in E. coli show differences in the temporal mode of processing in vivo. Cell. 1981;25(1):151-7. Epub 1981/07/01. PubMed PMID: 7023693.

43. Selkoe DJ. Folding proteins in fatal ways. Nature. 2003;426(6968):900-4. Epub 2003/12/20. doi: 10.1038/nature02264. PubMed PMID: 14685251.

44. Nakatogawa H, Ito K. The ribosomal exit tunnel functions as a discriminating gate. Cell. 2002;108(5):629-36. Epub 2002/03/15. PubMed PMID: 11893334.

45. Nakatogawa H, Ito K. Secretion monitor, SecM, undergoes self-translation arrest in the cytosol. Mol Cell. 2001;7(1):185-92. Epub 2001/02/15. PubMed PMID: 11172723.

46. Silhavy TJ, Shuman HA, Beckwith J, Schwartz M. Use of gene fusions to study outer membrane protein localization in Escherichia coli. Proc Natl Acad Sci U S A. 1977;74(12):5411-5. Epub 1977/12/01. PubMed PMID: 414221; PubMed Central PMCID: PMCPMC431741.

47. Lim B, Miyazaki R, Neher S, Siegele DA, Ito K, Walter P, et al. Heat shock transcription factor sigma32 co-opts the signal recognition particle to regulate protein homeostasis in E. coli. PLoS Biol. 2013;11(12):e1001735. Epub 2013/12/21. doi: 10.1371/journal.pbio.1001735. PubMed PMID: 24358019; PubMed Central PMCID: PMCPMC3866087.

48. Chatzi KE, Sardis MF, Tsirigotaki A, Koukaki M, Sostaric N, Konijnenberg A, et al. Preprotein mature domains contain translocase targeting signals that are essential for secretion. J Cell Biol. 2017;216(5):1357-69. Epub 2017/04/14. doi: 10.1083/jcb.201609022. PubMed PMID: 28404644; PubMed Central PMCID: PMCPMC5412566.

49. Derman AI, Puziss JW, Bassford PJ, Jr., Beckwith J. A signal sequence is not required for protein export in prlA mutants of Escherichia coli. Embo j. 1993;12(3):879-88. Epub 1993/03/01. PubMed PMID: 8458344; PubMed Central PMCID: PMCPMC413286.

50. Belogurov GA, Vassylyeva MN, Svetlov V, Klyuyev S, Grishin NV, Vassylyev DG, et al. Structural basis for converting a general transcription factor into an operon-specific virulence regulator. Mol Cell. 2007;26(1):117–29. doi: 10.1016/j.molcel.2007.02.021. PubMed PMID: 17434131; PubMed Central PMCID: PMCPMC3116145.

